# Regulation of defeat-induced social avoidance by medial amygdala DRD1 in male and female prairie voles

**DOI:** 10.1101/747923

**Authors:** Maria C. Tickerhoof, Luanne H. Hale, Adam S. Smith

**Author notes:** Corresponding author: Adam Smith, 1251 Wescoe Hall Drive, 5064 Malott Hall, Lawrence, KS, 66045.

## Abstract

Social interaction with unfamiliar individuals is necessary for species-preserving behaviors such as finding mates and establishing social groups. However, social conflict is a potential negative outcome to interaction with a stranger that can be distressing enough to cause an individual to later avoid interactions with other unfamiliar conspecifics. Unfortunately, stress research using a prominent model of social conflict, social defeat stress, has largely omitted female subjects. This has left a void in the literature regarding social strain on female stress biology and adequate comparison of the effect of sex in stress pathways. The prairie vole (*Microtus ochrogaster*) exhibits aggressive behavior in both sexes, making voles an attractive candidate to model social defeat in both sexes. This study sought to establish a model of social defeat stress in both male and female prairie voles, characterize behavioral changes in response to this stressor, and investigate the role of dopamine signaling in the response to social defeat stress. Defeated male and female prairie voles displayed social avoidance as well as an increase in expression of dopamine receptor D1 (DRD1) in the medial amygdala (MeA). Pharmacological manipulation of DRD1 signaling in the MeA revealed that increased DRD1 signaling is sufficient to induce a social avoidant state, and could be a necessary component in the defeat-induced social avoidance response. These findings provide the prairie vole as a model of social defeat in both sexes, and implicate the MeA in avoidance of unfamiliar conspecifics after a distressing social encounter.

## 1. INTRODUCTION

Social species have evolved to seek out interaction with unfamiliar individuals in order to thrive. Such interactions often have the end effects of establishing mating pairs or creating new social groups that work to defend each other from predators and share resources (O’Connell and Hofmann, 2011). Thus, interaction with unknown conspecifics can be highly advantageous. Yet, not all social interactions are positive in nature, and aggressive encounters are important for maintaining territorial boundaries and mate guarding. However, victims of aggressive encounters can become distrusting of other strangers in the future and thus avoid interactions with even non-aggressive conspecifics (Bjorkqvist, 2001; Toth and Neumann, 2013); this can have serious implications for the survival of an individual or even the species as a whole. This phenomenon applies to any social species, including more complex species such as humans, in which social conflict can act as a root cause for several mental illnesses that have social withdrawal as a primary symptom (Kendler et al., 2003). In order to understand and prevent the adverse effects social stress can have on an individual’s welfare, it is necessary to determine the effects social stress has on normal neurobiological function. Social defeat stress is an animal model of social conflict studied in a laboratory setting to examine the neurobiological underpinnings of deficits in social interaction (Toth and Neumann, 2013). Dopamine is a neurochemical that has been heavily implicated in the regulation of social avoidance following social defeat (Cao et al., 2010; Francis et al., 2015), as well as alterations to other behaviors attributed to defeat experience (Reguilon et al., 2017). Dopamine has also been implicated as having a role in humans suffering from mood disorders (Cervenka et al., 2012; Willner et al., 2005), supporting the validity of this model and the value of investigating the role of dopamine in stress pathology.

While social defeat stress research has been useful in determining the neurobiology of stress-induced maladaptive behavior, current modeling is largely restricted to males, mostly owing to the low incidence of female-directed behavior in most laboratory rodent species outside of maternal aggression (Bourke and Neigh, 2012). This has caused significant difficulty in characterization of neural systems governing social avoidance across both sexes, and has made it nearly impossible to make direct comparisons between males and females. A small number of studies have been performed to manipulate the behavior of male mice to attack females (Harris et al., 2018; Takahashi et al., 2017), though the concern of ethological validity is worth noting due to female-directed aggression being artificially induced. This concern is addressed through the use of species in which females naturally exhibit aggressive behavior towards each other, such as the California mouse (*Peromyscus californicus*) and Syrian hamster (*Mesocricetus auratus*) (Greenberg et al., 2015; Huhman et al., 2003). Studies using females of these species have implicated mesocorticolimbic dopamine signaling in the behavioral response to social defeat stress (Gray et al., 2015; Greenberg et al., 2015), similar to what has been observed in males. It has been observed that males and females of these species have variation aggressive behaviors directed toward the subject or differences in post-defeat behavioral phenotype (Huhman et al., 2003; Trainor et al., 2011; Trainor et al., 2013), allowing for the study of sex differences at the neurobiological level to explain the differences in behaviors and responsivity to conflict observed between males and females of those species. It is advantageous to explore the impact of conflict on the social dynamics and neurobiology processes in additional species in order to establish cross-species comparisons, as not all species display sex-specific responses to social conflicts.

Here, we use the prairie vole (*Microtus ochrogaster*) to investigate the effects of social defeat in both males and females. Following the formation of a pair bond, both male and female prairie voles display aggressive behavior toward unfamiliar conspecifics (Bowler et al., 2002); this aspect of prairie vole behavior can be utilized to subject animals of both sexes to social defeat without artificially inducing aggression, thus maintaining the appeal of defeat as a naturalistic stressor. Prairie voles also have a unique set of social behaviors significantly regulated by dopamine throughout the mesolimbic pathway and extended amygdala (Gobrogge and Wang, 2016; Northcutt and Lonstein, 2009). While the closely related Mandarin vole (*Microtus mandarinus*) has recently been used to model social defeat, a direct comparison of males and females has yet to be performed (Li et al., 2018; Wang et al., 2018). Additionally, recent work in developing transgenic prairie voles has begun to examine the effects of developmental gene manipulation on prairie vole sociality (Horie et al., 2018), increasing the utility of this species in examining the neurobiology connecting stress and social behavior as more tools become available. These reasons taken together present the prairie vole as an attractive model in further examining the role of dopamine in the impact of social defeat stress.

Although chronic defeat stress of ten days or more is well established to induce social avoidance and behavioral despair in mice (Golden et al., 2011) and Mandarin voles (Wang et al., 2018), we aimed to investigate whether a less severe defeat experience is still sufficient to impact normal prairie vole behavior, similar to what has been performed with the monogamous California mouse (Trainor et al., 2011; Trainor et al., 2013). The goal of this study was to establish a model of single and repeated social defeat in prairie voles of both sexes, and confirm the negative effect of social defeat on sociality in this species. In addition, because social defeat has not been characterized in prairie voles, we tested for changes in non-social behaviors as well to assist in understanding of this new model. Lastly, we aimed to determine defeat’s impact on dopamine receptor expression in key brain regions involved in regulation of sociality and stress response and their role in subsequent behavioral changes.

## 2. MATERIALS AND METHODS

### 2.1. Animals

Subjects were captive-bred sexually naïve male and female prairie voles descended from populations captured in southern Illinois. Voles were weaned on postnatal day 21 and housed with a same-sex conspecific in cages (29.2 L x 19.1 W x 12.7 H cm) containing corn cob bedding and crinkle paper nesting material with food and water *ad libitum*. Colony rooms were maintained on a 12L:12D photoperiod (lights on at 0600 hr) and at a temperature range of 21 ± 1°C. Resident aggressor voles were pair-bonded vasectomized male and female prairie voles that were previously screened to reach a minimum of three offensive attacks within five minutes of intruder presentation. All behavioral testing was performed between 0900-1600hr. All procedures were conducted in accordance with the National Institutes of Health Guide for the Care and Use of Laboratory Animals and the Institutional Animal Care and Use Committee at the University of Kansas.

### 2.2. Behavior

All behavior tests were recorded with video and manually scored using JWatcher (UCLA).

#### 2.2.1. Defeat procedure

The social defeat protocol was adapted from a previous resident-intruder method published using female prairie voles (Smith et al., 2013). At least one day prior to defeat, resident aggressor pairs were moved into large cages (47.6 L x 26.0 W x 15.2 H cm) to establish home cage territory. On days of social defeat, aggressors and subjects were transported to our behavioral testing facility and allowed to acclimate to the environment for at least 30 min. During the acclimation period, the aggressors’ mates, nesting material, and hopper containing food and water were removed from the cage. At the start of social defeat conditioning, the subject vole (i.e. the intruder) was placed into the aggressor’s (i.e. resident’s) cage and exposed to a 15-min period of physical. Following this 15-min session, the intruder vole was placed into a stainless-steel corral (9 L x 9 W x 14 H cm) in the center of the resident’s cage for a 45-min threat session for continued exposure to visual and olfactory cues from the resident. At the end of the 45-min threat session, every defeated vole was examined for injuries then returned to the home cage. Although rare, any animal that received significant injury was immediately discontinued in defeat exposure and removed from the study as described in an established chronic defeat protocol using mice (Golden et al., 2011).

*Control* voles were placed into an empty novel cage for one hour, then returned to the home cage, and repeated this brief separation for all three days. *Single defeat* voles received control treatment for two days, then were defeated on the third. *Repeated defeat* voles were defeated all three days, by a different resident each day.

#### 2.2.2. Home Cage Behavior

In-group dynamics were evaluated during 15-min sessions before conditioning (*baseline*), during return to the home cage after the final conditioning period (*reactivity*), and six days after the end of conditioning (*recovery*). For baseline recording, the cage contents (i.e., nesting material, food, and water) were removed and the two animals were allowed to habituate for 15 minutes before beginning recording. For the other two recording sessions, cage contents were removed from the home cage 15 minutes prior to the end of the subject’s conditioning to allow the cagemate to habituate to the empty cage before the return of the subject. The subject vole was observed for the following behaviors: proximity with the cagemate (defined as distance less than one-half body length), sniffing of the cagemate, grooming of the cagemate (allogrooming), and stationary side-by-side contact. The Hinde index, a measure of which individual is responsible for maintaining contact in a dyadic interaction (Silk et al., 2013), was also calculated.

#### 2.2.3. Social Preference/Avoidance Test

The social preference/avoidance (SPA) test is adapted from the three-chamber social approach test (Toth and Neumann, 2013). The subject vole was placed into the center of a three-chamber arena (75 L x 20 W x 25 H cm) containing a thin layer of corn cob bedding and permitted to habituate to the full arena for 10 min or until the subject enters each chamber at least once, whichever comes last. Following this habituation period, subjects were sequestered into the center chamber and stimuli in stainless steel corrals were placed against the far walls of opposite chambers in a counterbalanced manner. Stimuli were composed of a novel same-sex prairie vole (social) and a novel black rubber stopper (object). After stimuli were placed, the barriers separating the center chamber from the rest of the arena were removed, and the subject vole was permitted to explore the arena and stimuli for 10 min. Active investigation (sniffing) of each stimulus was analyzed.

#### 2.2.4. Elevated Plus Maze

The elevated plus maze (EPM) was comprised of two open arms (36.5 L×7 W cm) at approximately 420 lux and two closed arms (36.5 L×7 W×20 H cm) at approximately 20 lux that cross in the middle, and is elevated 60 cm off the ground. Subjects were placed in the center of the arena facing an open arm, then recorded for a period of 5 minutes. The number of arm crosses and time spent in the open arms were analyzed.

#### 2.2.5. Forced Swim Test

A plexiglass cylinder (20 W x 30 H cm) was filled with 4 L of tap water at 24° ± 1° C. The subject was placed into the cylinder and left to swim for 5 min. Immobility and swimming/struggling behaviors were analyzed.

#### 2.2.6. Open Field Test

Subject voles were placed into the center of a force plate actometer (42 L x 42 W cm) (Fowler et al., 2001) and permitted to explore the arena for 30 min. Time spent in the central portion of the arena and total locomotion were analyzed.

#### 2.2.7. Sucrose Preference Test (SPT)

Two 50 mL water bottles were placed into the hopper of subjects’ home cages, one containing distilled water and the other containing a 0.25% solution of sucrose in water. This concentration of sucrose was chosen after an optimization pilot experiment as 1% concentration previously used for prairie voles (Grippo et al., 2007) led to over-consumption of the sucrose solution in our colony. The two bottles remained in the home cage hopper for 3 days, then subjects were individually housed for a 4-day testing period. The bottles were switched each day to avoid side preference. Volume of each bottle was measured at the beginning and end of each day to determine consumption.

### 2.3. Biological characterization

#### 2.3.1. Western Blots

Brains were sectioned with a cryostat at 150 microns, and bilateral tissue punches containing the prefrontal cortex (PFC), nucleus accumbens (NAcc), basolateral amygdala (BLA), central amygdala (CeA), and medial amygdala (MeA) were collected. Tissue punches were lysed in RIPA buffer and the protein extract stored at −80° C until analysis. Total protein concentration was determined using DC assay, and protein extracts containing 15μg of total protein were loaded into 15% Tris-HCl gels. Following electrophoresis and transfer, membranes were stained with 5% Ponceau S as a measure of total protein content (Thacker et al., 2016). Ponceau S was rinsed away with T-TBS and membranes were blocked with 5% nonfat milk in T-TBS. Membranes were incubated with anti-dopamine receptor 1 (DRD1) or 2 (DRD2) primary antibody (**Table S2**) in 5% nonfat milk in T-TBS overnight at 4°C. Following primary incubation, membranes were exposed to appropriate HRP-conjugated secondary antibody (**Table S2**), scanned, and bands quantified with ImageJ (NIH).

#### 2.3.2. Corticosterone ELISA

Trunk blood was collected into EDTA-coated tubes and plasma was separated by two centrifugation cycles at 2000xG, then stored at −80°C until analysis. A 1:1000 dilution of plasma in assay buffer was analyzed with a corticosterone ELISA kit (Cayman Chemical, Cat #501320) following manufacturer instructions. Samples were run in duplicate on a single plate. All sample CVs were <10%, and intra-assay CV was 3.48%.

### 2.4. Pharmacological manipulation of MeA DRD1

#### 2.4.1. Stereotactic surgery

Voles were deeply anesthetized with an i.p. injection of a ketamine/dexmedetomidine cocktail (75/1 mg/kg), then head-fixed into a stereotax (Stoelting). Guide cannula (26 gauge, PlasticsOne) were implanted bilaterally aimed toward the MeA (5° angle, A/P −1.05 M/L ±2.91 D/V −4.82) and affixed to the skull with two surgical screws, ethyl-2-cyanoacrylate-based adhesive, and dental cement. Anesthesia was reversed with atipamezole (2.5 mg/kg, intraperitoneal) and voles were returned to a new cage with larger dimensions to accommodate for the implants (36.2 L x 20.3 W x 14.0 H cm).

#### 2.4.2. Drug preparation

The drugs used were a DRD1-selective agonist (SKF 38393, Tocris Bioscience, Cat #09-221-00) and DRD1-selective antagonist (SCH 23390, Millipore, Cat #505723). Drugs were prepared ahead of time by dissolving drug in ddH_2_O at a 10X concentration (20μg/mL SKF 38393, 200μg/mL SCH 23390) and freezing at −20°C for no longer than one month. On the day of testing, drug stock was diluted to a 1X concentration using artificial cerebrospinal fluid (aCSF; 150mM Na^+^, 3.0 mM K^+^, 1.4 mM Ca^2+^, 0.8 mM Mg^2+^, 1.0 mM P^3-^, 155 mM Cl^−^).

#### 2.4.3. Site-specific drug administration

Thirty minutes before SPA, 200nL aCSF containing no drug (CSF), 0.4 ng SKF 38393 (SKF), or 4 ng SCH 23390 (SCH), was injected bilaterally with a 1mm projection using 33 gauge internal cannula (PlasticsOne) into the MeA through the guide cannulae. The rate of infusion (200 nl/min) was controlled with an infusion pump (KD Scientific). The subjects were returned to the home cage for 20 min, then placed in the empty testing arena for a 10-min habituation prior to testing for a total 30-min drug activation time. These doses and time period were selected based on previous literature using these drugs in prairie voles (Liu et al., 2011).

### 2.5. Experimental design

The timeline for each behavioral experiment is presented in **Figure S1.**

#### 2.5.1. Experiment 1: Effects of single and repeated defeat on behavior and neurobiology

Since examination of alterations in home cage interaction was a goal of this first experiment, only one vole in each cage received conditioning (control, single, or repeated defeat) and the cagemate was left un-manipulated. 69 total animals (Control: 11M/11F, single defeat: 11M/12F, repeated defeat: 12M/12F) were used in this experiment. Home cage interaction with the cagemate was examined as described in *2.2.2*. Starting one week after conditioning, subjects were tested with SPA, EPM, and FS, separated a week apart. On the third week, immediately after FS, animals were euthanized and brains were harvested, flash frozen on dry ice, then stored at −80° C until processing.

One animal each in the male single defeat and female repeated defeat groups were excluded due to insufficient aggression, leading to a final sample size of 10-12 per group. One “reactivity” home cage recording, 11 “recovery” home cage recordings, and 1 FS recording were lost due to technical difficulties.

#### 2.5.2. Experiment 2: Further characterization of behavior with both cagemates in same condition

Because home cage observation was not incorporated in future experimental designs, in the interest of reducing the number of animals used we aimed to determine whether conditioning both cagemates would impact subsequent behavior. Both animals in each cage received the same conditioning (control or repeated defeat). 36 total animals (Control: 8M/8F, repeated defeat: 12M/8F) were used in this experiment. Starting one week after conditioning, subjects were tested with SPA, EPM, FS, OFT, and SPT. Brain tissue was not analyzed due to the confound of social isolation at the end of the experiment for the four-day SPT protocol.

Two repeated defeat males were removed from the study due to injury, leading to a final sample size of 8-10 per group. Two repeated defeat males were removed after SPA and two control females were removed after FS due to unexpected complications unrelated to the defeat protocol.

#### 2.5.3. Experiment 3: Pharmacological manipulation of social interaction in SPA

40 total animals (Control-CSF: 10, Control-SKF: 10, Defeat-CSF: 10, Defeat-SCH: 10) were used in this experiment. Sex was not an experimental factor and thus males and females were evenly distributed. Stereotactic surgery was performed at least one week prior to conditioning. Both animals in each cage received the same conditioning (control or repeated defeat), but may not necessarily have received the same drug treatment. One week after conditioning, subjects received drug treatment and 30 minutes later were tested in SPA as described in *2.4.3*. Immediately after SPA, animals were euthanized via rapid decapitation and trunk blood was collected for corticosterone analysis. Brains were also harvested for confirmation of cannulae placement.

Animals determined to have incorrect cannula placement (Control-CSF: 2, Control-SKF: 3, Defeat-CSF: 2, Defeat-SCH: 1) were excluded from analysis, leading to a final sample size of 7-9 per group.

### 2.6. Data analysis

#### 2.6.1. Statistical analysis

Statistical outliers, defined as data points two standard deviations above or below the mean, were excluded from analysis. Data points excluded due to being statistical outliers were removed from the final sample sizes as indicated in *2.5.1-3*.

RIT behavior was analyzed with repeated measures ANOVA (between-subjects: sex, within subjects: social status, day). All behavioral and western blot data in Experiments 1 and 2 were analyzed with a two-way ANOVA (between-subjects: sex, condition), with the exception of SPA and SPT which were a three-way repeated measures ANOVA (included within-subjects: stimulus type) and analysis of the CeA which was performed with Kruskal-Wallis due to lack of normality. Experiment 3 corticosterone was analyzed with one-way ANOVA (between-subjects: defeat/drug group) and SPA with repeated measures ANOVA (included within-subjects: stimulus type). Pairwise comparisons were performed with Student-Neuman-Keuls (SNK) post-hoc analysis. Effects were considered significant at p<0.05. All data are presented as Mean ± SEM.

## 3. RESULTS

### 3.1. Characterization of social defeat in prairie voles of both sexes

Only one publication has studied social defeat in prairie voles (Smith et al., 2013), and this was limited to a single exposure in females. To first elucidate if there are differences between male and female prairie voles during social defeat, offensive and defensive behaviors of residents and intruders of both sexes were compared across three days. Between all three days, residents overall displayed significantly more total (both physical and non-physical) offensive behaviors than intruders (F[_1,14_]=107.460, p<0.001; **Figure 1A**), and intruders displayed more defensive behaviors than residents (F[_1,14_]=139.108, p<0.001; **Figure 1C**). A breakdown of the average occurrence of individual behaviors (**Tables S1 and S3**) revealed that, with the exception of upright posturing, residents displayed each offensive behavior significantly more than intruders (all F[_1,14_]>10.337, all p<0.01; **Figure 1B**). Similarly, each individual defensive behavior was displayed almost exclusively by intruders (all F[_1,14_]>5.576, all p<0.05; **Figure 1D**). The only sex differences observed were a slightly lower average number of physical attack by female residents (F[_1,14_]=7.764, p<0.05), as well a higher rate of retaliatory attack by male intruders than females (F[_1,14_]=4.921, p<0.05), though this behavior was rarely observed. Altogether, these results indicate that male and female prairie voles exhibit nearly identical aggression in the resident-intruder paradigm, thus confirming that subject voles of both sexes received similar defeat experiences.

**Figure 1:**
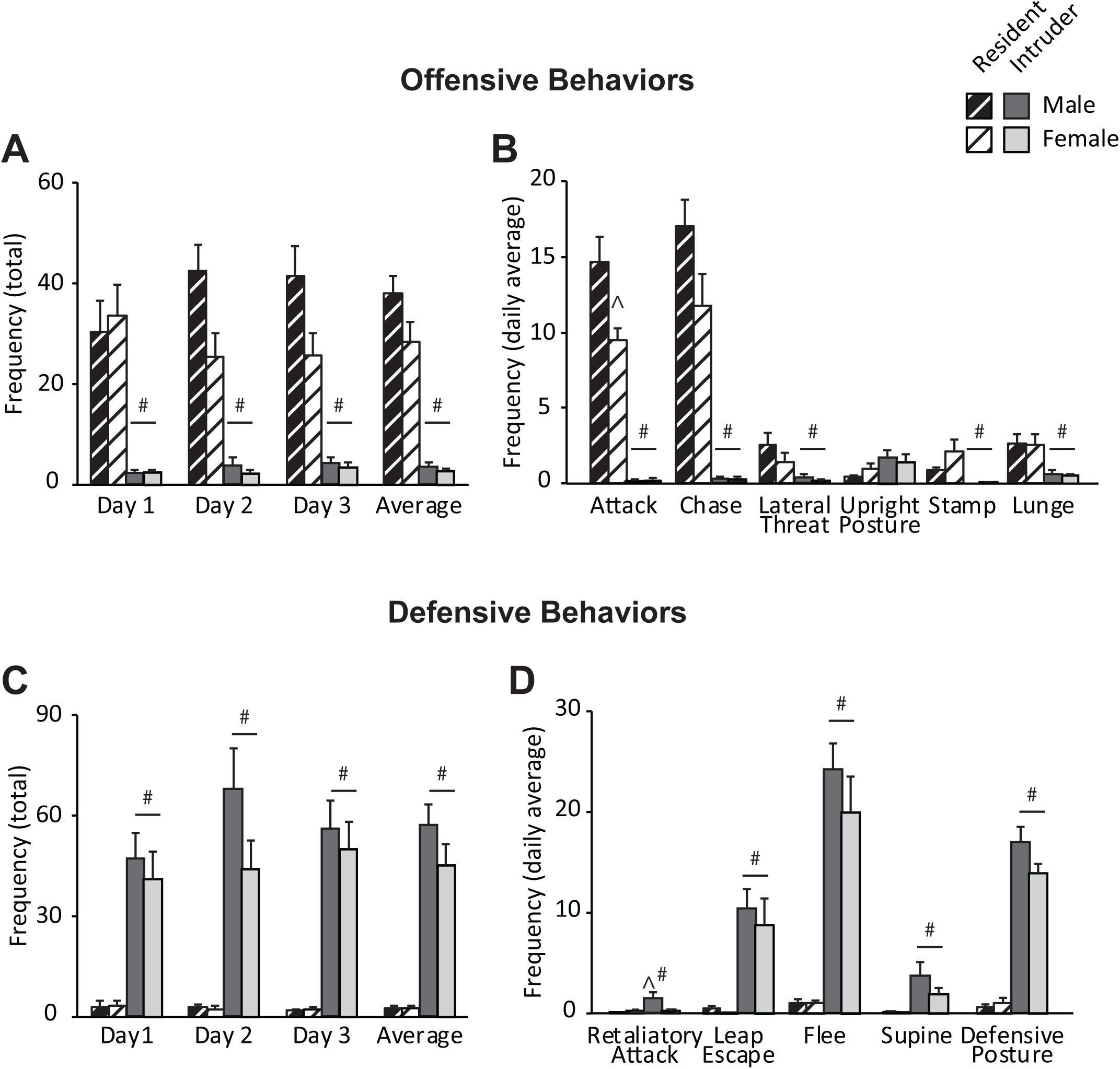
Male and female prairie voles receive similar defeat experiences and display nearly identical defensive behavior during defeat. **A)** Residents of both sexes display significantly more total offensive behaviors than defeated intruders, **B)** which applies to almost all individual behavior types. **B)** Defeated intruders of both sexes are nearly exclusively responsible for the display of total defensive behaviors during defeat, **D)** including every individual behavior type. Definitions of individual behaviors are presented in **Table S1**. *(n=8 per group)* #p<0.05 vs resident, ^p<0.05 vs other sex.

### 3.2. Effects of social defeat on behavioral phenotype

#### 3.2.1. Home cage interaction

To investigate changes in in-group social interaction, behavior between Experiment 1 subjects and their cagemates were examined. Voles of both sexes designated for all conditions did not differ in any observed behavior at baseline observation (data not shown). Immediately following the final session of conditioning (**Figures 2A-C**), voles that had experienced repeated defeat sniffed the cagemate more than controls (F[_2,56_]=5.095, p<0.01), and both single and repeated defeat voles groomed the cagemate significantly less than controls (F[_2,56_]=3.881, p<0.05). Omnibus two-way ANOVA revealed a significant sex x condition interaction for side-by-side contact (F[_2,56_]=3.401, p<0.05), but this effect failed to be clarified with post-hoc analysis (data not shown). Animals were also analyzed for changes in Hinde index, a measure of which individual in a dyadic interaction is responsible for maintaining contact (Silk et al., 2013). In the reactivity observation period, there was a main effect of condition such that defeated voles had a positive Hinde index that was significantly higher than that of controls (F[_2,56_]=14.781, p<0.001), indicating that the defeated animal was more responsible for maintaining contact with the cagemate. All significant effects in social behavior and Hinde index were lost by the recovery observation period (**Figures 2D-F**), suggesting that acute changes in in-group social dynamics had normalized by the time other behavioral analyses were performed.

**Figure 2:**
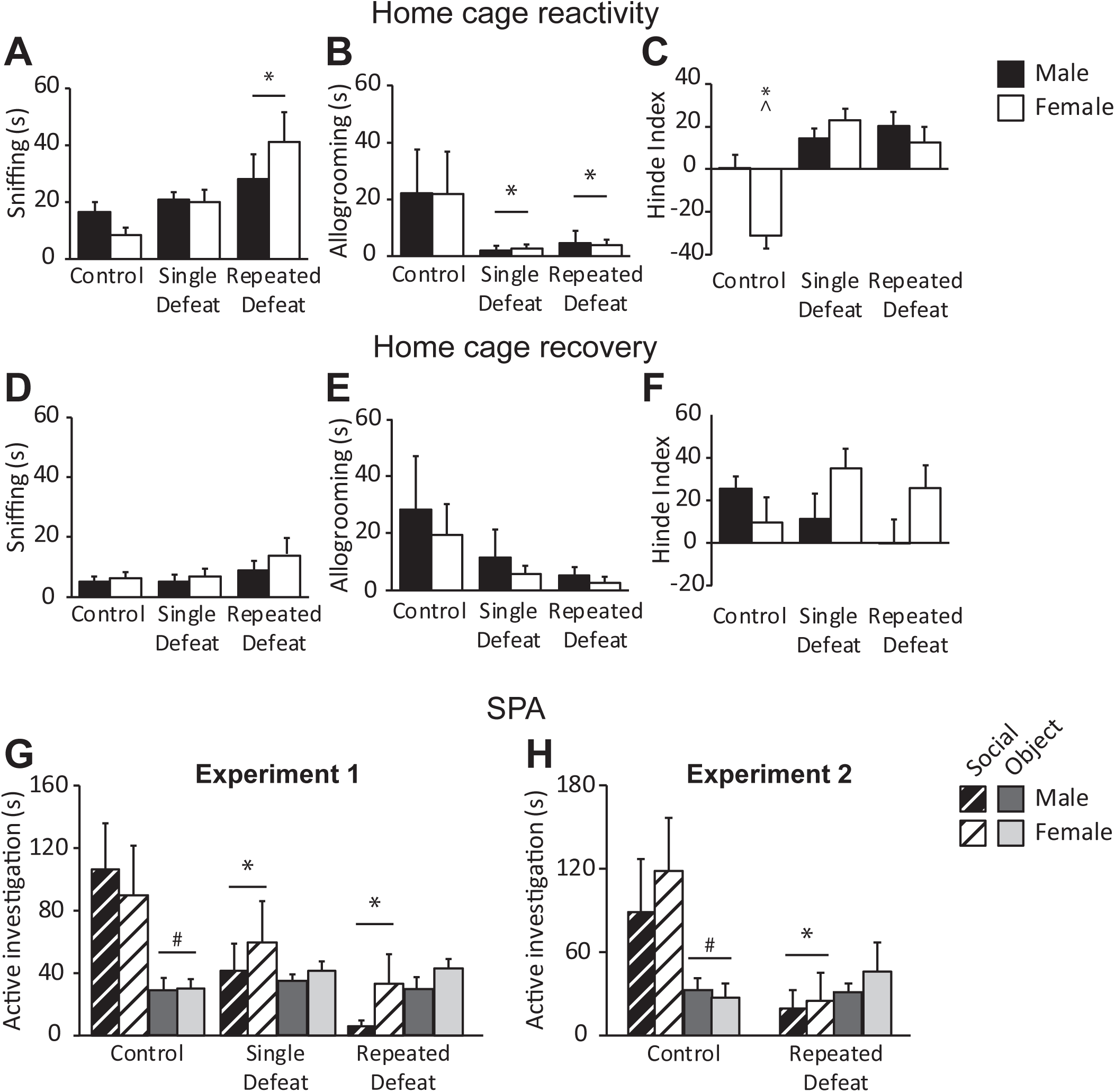
Social defeat impacts interaction with the cagemate acutely but not long-term, and significantly impairs investigation of a novel conspecific. **A-C)** Sniffing and allogrooming of the cagemate by defeated animals, as well as Hinde index, are different from their non-defeated counterparts immediately following the final session of conditioning. *(n=7-12 per group)* **D-F)** Behaviors in the home cage interaction six days after the final session of conditioning are no longer different between controls and defeated voles. *(n=7-10 per group)* **G)** Social defeat stress significantly impairs investigation of an unfamiliar conspecific in both sexes both when the cagemate is un-manipulated *(n=9-12 per group)* **H)** and when both cagemates receive the same conditioning. *(n=7-8 per group)* *p<0.05 vs control, #p<0.05 vs social stimulus, ^p<0.05 vs other sex. *SPA-Social preference/avoidance test*.

#### 3.2.2. SPA

To investigate changes in social interaction with an unfamiliar conspecific, social behavior of subject voles was examined with the SPA test. In Experiment 1, defeat conditioning significantly decreased social approach without affecting novel object investigation (F[_2,56_]=7.094, p<0.01; **Figure 2G**). This effect was independently replicated in Experiment 2 (F[_1,26_]=13.371, p<0.001; **Figure 2H**), indicating that similarly conditioning cagemates had no effect on subsequent behavioral phenotype. No effect of sex on investigation time for any condition was observed. These results suggest that social defeat stress induces a social avoidance phenotype in both male and female prairie voles.

#### 3.2.3. EPM

In Experiment 1, there was no effect of defeat on frequency of crosses into the open arm (**Figure 3A**). However, defeated voles spent less time in the open arm (F[_2,52_]=3.139, p<0.05; **Figure 3A**) and trended towards an increased latency to enter the open arm (F[_2,52_]=3.037, p=0.06; **Figure 3B**). Similar effects were also seen in Experiment 2, with the increased open arm latency reaching significance (Open arm duration: F[_1,23_]=6.343, p<0.05; Open arm latency: F[_1,23_]=5.954, p<0.05; **Figures 3D-E**). While females overall spent more time in the open arms in Experiment 2, there was no interaction between sex and condition in either experiment. Therefore, social defeat stress altered behavior in the elevated plus maze in a manner indicative of a potentially anxious state in voles of both sexes.

**Figure 3:**
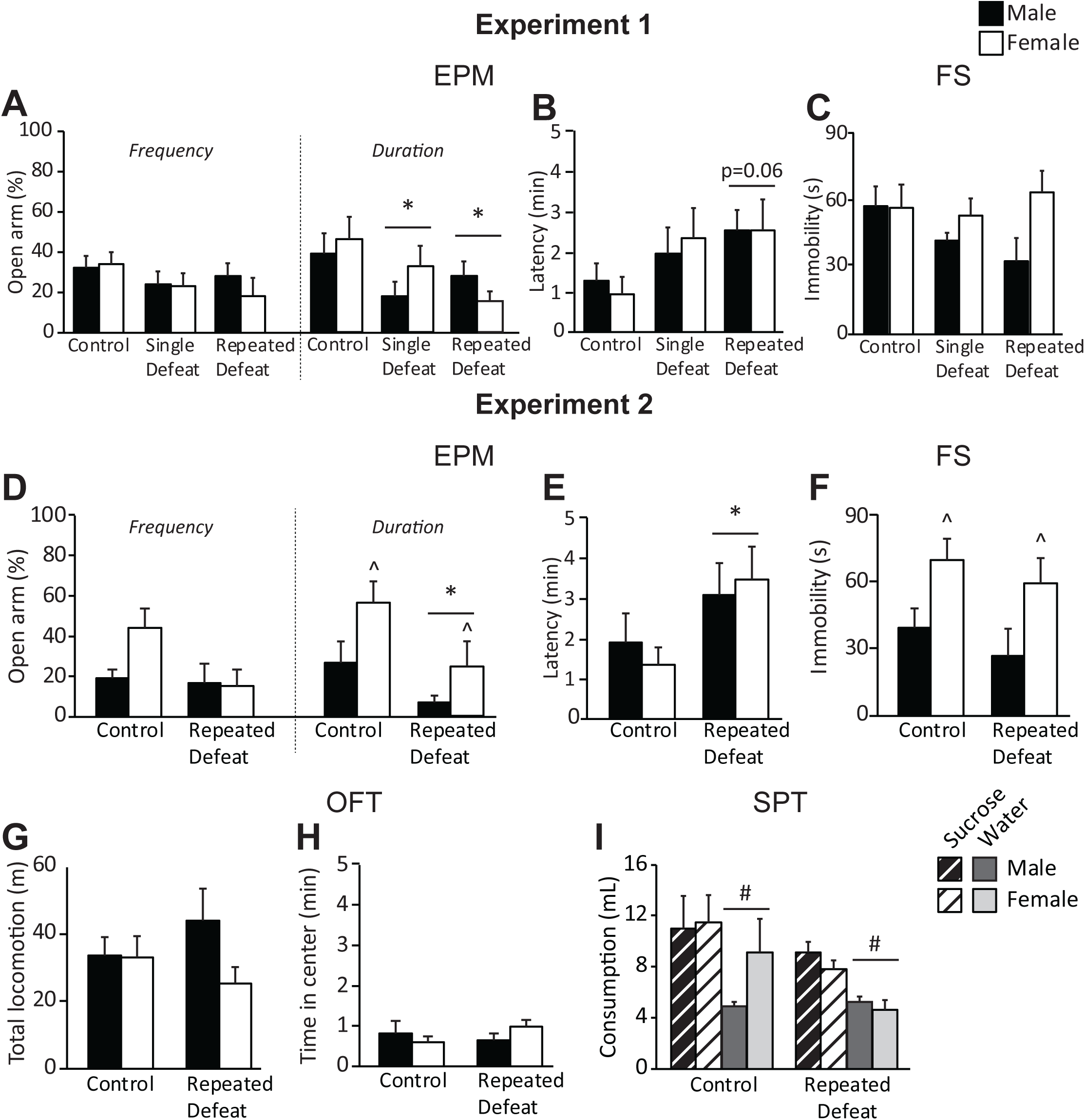
Open arm exploration in EPM is impaired in defeated animals, but defeat did not impact other non-social behavioral measures. **A-B)** Open arm exploration in EPM is impaired in defeated voles of both sexes as measured by duration of time spent in the open arm and latency to first open arm entry. *(8-11 per group)* **C)** Social defeat experience did not impact passive coping behavior as measured by immobility in FS. *(n=10-11 per group)* **D-E)** Voles display similar behavioral abnormalities in EPM when both cagemates are defeated as when only one animal is defeated. *(n=6-8 per group)* **F)** Both animals receiving similar conditioning still led to no effect of defeat in FS. *(n=6-8 per group)* **G-H)** Social defeat did not impact time spent in the center of the arena nor total locomotion in the OFT. *(n=6-8 per group)* **I)** Both control and defeated voles exhibited a significant preference for the 0.25% sucrose solution over the plain water. *(n=6-8 per group)* *p<0.05 vs control, #p<0.05 vs sucrose solution, ^p<0.05 vs other sex. *EPM-Elevated plus maze test; FS-Forced swim test; OFT-Open field test; SPT-Sucrose preference test*.

#### 3.2.4. FS

There was no main effect of condition on time spent immobile in FS for either experiment (**Figures 3C, 3F**). There was a main effect of sex in Experiment 2 such that females overall spent more time immobile than males (F[_1,24_]=9.195, p<0.01), but there was no effect of sex in Experiment 1 nor an interaction between sex and condition in either experiment. This suggests coping strategies are not impacted by this defeat protocol in prairie voles.

#### 3.2.5. OFT

There was no effect of sex or defeat on time spent in the center of the open field or on total locomotion over the course of the test (**Figures 3G-H**).

#### 3.2.6. SPT

While there was an effect of defeat such that defeated voles of both sexes drank less overall than control voles (F[_1,22_]=5.447, p<0.05), both control and defeated voles had a significant preference for the sucrose solution over the regular water (F[_1,22_]=14.060, p<0.001; **Figure 3I**). Therefore, social defeat experience had no effect on anhedonia in prairie voles of either sex.

### 3.3. Evaluation of dopamine receptors following defeat

To examine the effect of defeat on dopamine systems, DRD1 and DRD2 protein expression was characterized in several key regions implicated in social behavior and stress reactivity. Three weeks after defeat experience, there were no changes in DRD1 or DRD2 expression in the NAcc, PFC, or BLA (**Table 1**). In the CeA there was an interaction between sex and condition for DRD1 expression (H_5_=15.238, p<0.01; **Table 1**), such that repeated defeat female voles had a decrease in DRD1 expression relative to both females and males in all other conditions. In the MeA, there was a main effect of condition for DRD1 expression, and post-hoc analysis determined repeated defeat voles had an increase in levels of DRD1 relative to control and single defeat voles (F[_2,53_]=3.550, p<0.05; **Figure 4**). There were no changes in DRD2 expression in the CeA or MeA. These results suggest an effect of defeat on amygdalar dopamine receptor expression, and implicate a role of amygdalar dopamine signaling in the behavioral response to social defeat stress.

**Table 1:**
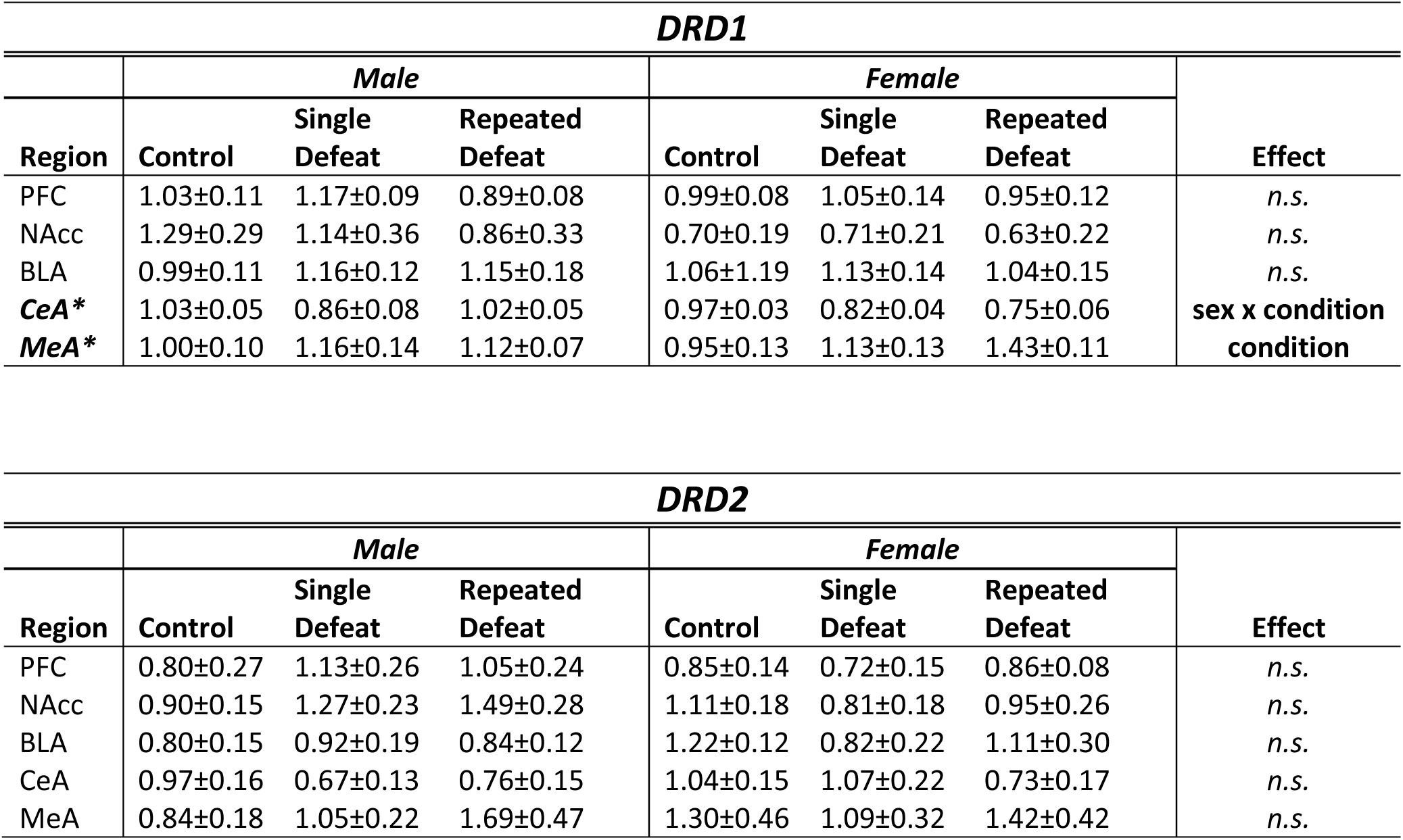
Western blot analysis of dopamine receptor expression in mesocorticolimbic and extended amygdala regions. *p<0.05 *PFC-Prefrontal cortex; NAcc-Nucleus accumbens; BLA- Basolateral amygdala; CeA- Central amygdala; MeA- Medial amygdala*.

**Figure 4:**
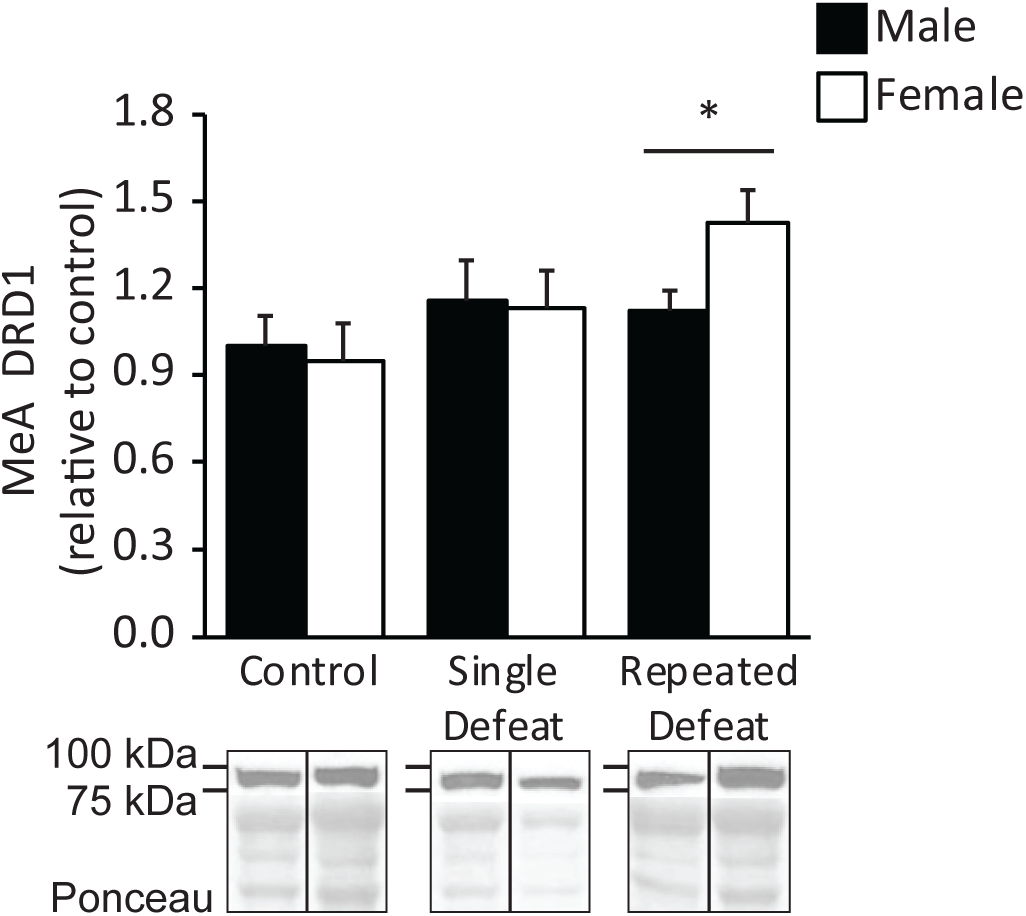
DRD1 expression in the MeA is increased in defeated prairie voles. Voles in the repeated defeat group had an increased expression of DRD1 in the MeA relative to controls. *(n=9-10 per group)* Representative bands displayed were selected due to an optical density/Ponceau ratio near average for their respective group. *p<0.05 vs control. *DRD1-Dopamine receptor subtype 1; MeA-Medial amygdala*.

### 3.4. Role of medial amygdala DRD1 in avoidance of novel conspecifics

In order to confirm the role of MeA DRD1 in the social avoidance behavior following social defeat stress, prairie voles of both sexes were implanted with guide cannulae aimed for the MeA to directly administer DRD1 drugs to this region (**Figure 5A**). To emulate the effects of increased DRD1 activity, control voles either received vehicle or 0.4ng SKF 38393, a DRD1-selective agonist. Conversely, repeated defeat voles received either vehicle or 4ng SCH 23390, a DRD1-selective antagonist. Due to the lack of effect of sex in both SPA behavior and MeA DRD1 expression, males and females were distributed evenly across all groups.

**Figure 5:**
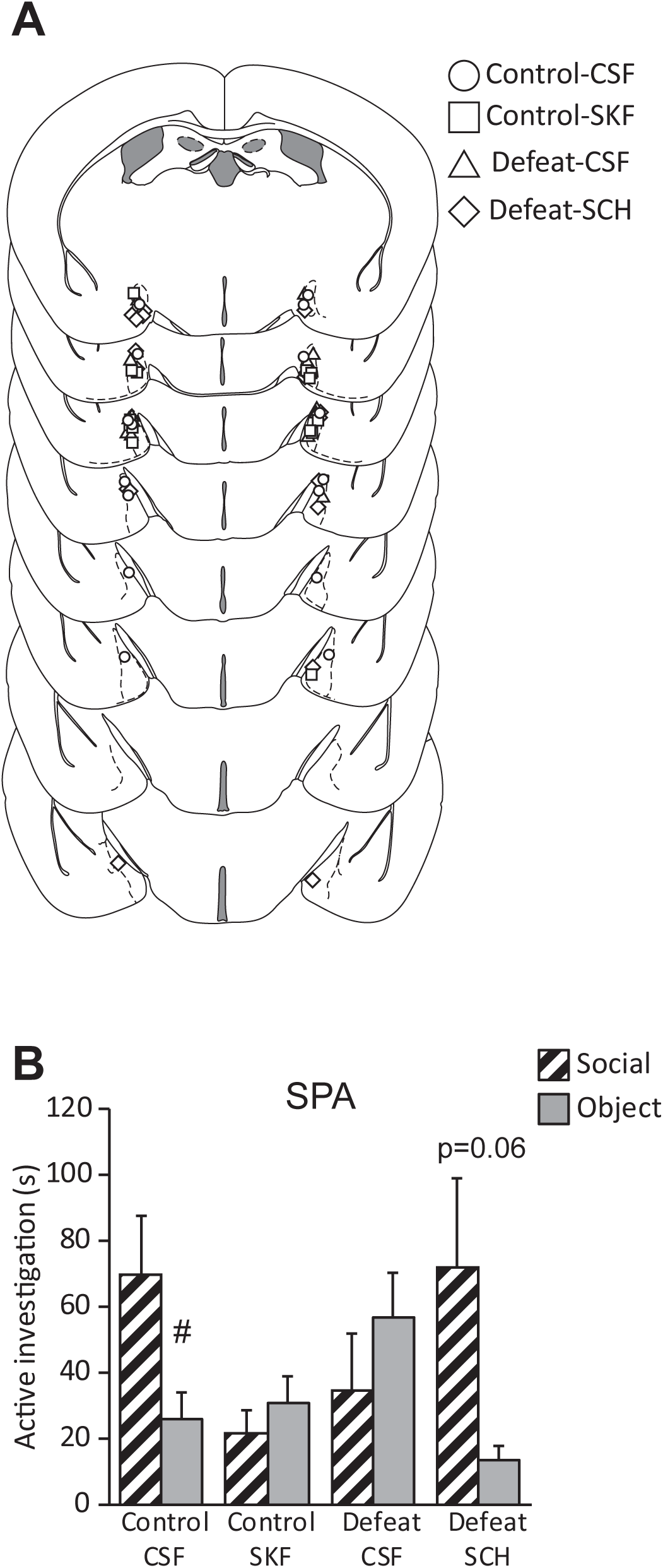
Increased MeA DRD1 activity is sufficient to induce social avoidance without defeat experience and could be necessary for the social avoidance response following defeat. **A)** Distribution of successful stereotactic hits included in data analysis. Brain schematics adapted from Franklin & Paxinos Mouse Brain Atlas (3ed. compact) **B)** Site-specific administration of the DRD1 agonist SKF 38393 into the MeA of stress-naïve voles led to social avoidance in SPA. Administration of the DRD1 antagonist SCH 23390 into the MeA of defeated voles nearly restored a social preference. #p<0.05 vs social stimulus. *SPA-Social preference/avoidance test; DRD1-Dopamine receptor subtype 1; MeA-Medial amygdala*.

#### 3.4.1. SPA

Following treatment with a DRD1-selective drug, there was a significant interaction between defeat/drug group and stimulus investigation (F[_3,26_]=3.113, p<0.05; **Figure 5B**). As observed in previous experiments, control voles that received vehicle treatment before the SPA test exhibited a robust preference for investigation of the novel vole over the object stimulus. Control voles that received administration of the DRD1 agonist SKF 38393 restricted social investigation to the extent that no preference between the social and object stimuli was observed. This behavioral pattern mirrored the vehicle-treated defeated voles, which also displayed social avoidant behavior as previously observed. Defeated voles that received a dose of the DRD1 antagonist SCH 23390 trended towards preferring the social stimulus over the object stimulus. These results indicate that increased DRD1 signaling in the MeA is sufficient to induce a social avoidance phenotype without defeat experience, and may be a necessary component of the behavioral response following social defeat stress.

#### 3.4.2. Corticosterone

There was no effect of defeat/drug group on plasma corticosterone levels (Control-CSF: 557.9±84.6 ng/mL; Control-SKF: 549.2±65.9 ng/mL; Defeat-CSF: 545.1±63.1 ng/mL; Defeat-SCH: 667.5±50.2 ng/mL).

## 4. DISCUSSION

This study was designed to characterize the behavioral effects of social defeat stress in prairie voles of both sexes. First, we confirmed that voles intruding on the territory of a pair-bonded resident will be subsequently defeated in an aggressive encounter, and that voles of both sexes received nearly identical defeat experience. In addition, we determined a single or repeated defeat experience induces a social avoidance phenotype in both male and female prairie voles. This is consistent with chronic defeat models using male mice and rats (Toth and Neumann, 2013). The defeat-induced social avoidance phenotype is sometimes exclusively referred to as a “depressive-like” phenotype, with the interpretation that an avoidance of social interaction is due to decreased motivation. However, the other behavioral tests performed in this study do not support that conclusion in our model. The lack of effect of defeat in FS and SPT not only indicate that defeated voles did not exhibit a changes in coping strategy or anhedonia, but also suggest that the social avoidance phenotype is not due to a lack of motivation, since active coping behaviors and motivation to drink sweetened water did not differ from controls. Due to the design of the experimental timeline, there may be the initial concern that the defeat-induced phenotype does not persist the full five weeks that behavioral measures were taken; however, our lab has found that the social avoidance phenotype following social defeat stress persists at least eight weeks in both male and female prairie voles (L.H. Hale, M.C. Tickerhoof, and A.S. Smith, unpublished observations). More likely, the lack of effect in these measures may be due to the defeat protocol used, as chronic defeat (10-14 consecutive days) in mice has been shown to impact behavior in the forced swim and sucrose preference tests, but sub-chronic defeat (3 consecutive days) does not (Francis et al., 2015). Notably, 14 days of social defeat in females of the closely-related Mandarin vole does indeed lead to increased immobility in FS and tail suspension tests (Wang et al., 2018). Whether chronic defeat stress would significantly impact behavioral phenotype in prairie voles beyond what is observed in the current single and repeated defeat model may warrant further investigation.

Interestingly, we noted increased social interaction with a familiar cagemate immediately upon return to the home cage from the final session of conditioning, perhaps as an attempted stress coping strategy as social buffering has been reported in pair bonded voles (Burkett et al., 2016; Smith and Wang, 2014). The presence of an unstressed cagemate did not appear to diminish the negative effects of defeat, however, since the effect of defeat was robust and replicable regardless of if the cagemate was also defeated. This is likely because this study used same-sex sexually naïve prairie voles, thus suggesting that the unique nature of the pair bond is necessary for effective social buffering, but the nature of the stressor and experimental timeline could also play a role. One week after social defeat, voles displayed rather normal social interaction with their cagemate but were avoidant of contact with a novel, non-aggressive conspecific, indicating that the social avoidance phenotype is exclusive to unfamiliar conspecifics, and may not significantly impact in-group social dynamics.

The null effects in the OFT and corticosterone assay may be due to limitations of the model itself. Multiple studies measuring behavior in the open field test with prairie voles have shown that prairie voles perform poorly in this test with low levels of time spent in the center even without experimental manipulation (Kenkel et al., 2014; Lieberwirth et al., 2012), and thus it may more appropriate as solely a measure of total locomotion in this species. The levels of blood corticosterone measured in this study were also consistent with baseline corticosterone levels in other prairie vole studies (McNeal et al., 2017; Taymans et al., 1997), indicating that peripheral stress markers do not appear to be affected following the SPA test regardless of defeat status or drug treatment. Social defeat does acutely increase blood corticosterone levels in female prairie voles (Smith et al., 2013) and Mandarin voles (Li et al., 2018); however, baseline corticosterone may not be impacted chronically by social defeat, even when later presented with a novel social stimulus. This finding appears to be consistent with data observed in California mice, where no changes in blood corticosterone are observed following a social interaction test four weeks after defeat experience (Trainor et al., 2011). An additional possible explanation is that since avoidant animals were able to separate themselves from the social stimulus by 75cm, the HPA axis may not have responded to the source of stress because the act of avoidance in itself acts to diminish the physiological response to a source of stress (Smith et al., 2013).

The discovery that social defeat affects protein expression in MeA, which then subsequently regulates defeat-induced behavioral changes, may coincide with, or emerge beyond, the well-established role of this region in responding to odor-dependent unconditioned threats. The MeA is part of the extended olfactory system, receiving projections from both the main and accessory olfactory bulbs (Cadiz-Moretti et al., 2016) and regulates discrimination between conspecific and heterospecific (such as predator) odors (Meredith and Westberry, 2004). Thus, the MeA regulates innate behaviors that bring together environmental information and social cues to promote survival responses. Activity in the MeA increases in response to exposure to predator odor (Dielenberg et al., 2001; McGregor et al., 2004), and inhibition of MeA activity reduces fear-related behaviors in response to predator odors (Li et al., 2004; Müller and Fendt, 2006). The MeA is also activated in response to illness-related odors, and inhibition of neuropeptidic receptors in this region reduce avoidance of sick conspecifics (Arakawa et al., 2010). MeA neurons are activated after aggressive encounters (Choi et al., 2005), and neurons in the MeA project to regions of the bed nucleus of the stria terminalis (BNST) and hypothalamus responsible for defensive and avoidance behaviors (Choi et al., 2005; Miller et al., 2019; Pardo-Bellver et al., 2012). Recent literature would suggest that this is at least partially mediated by DRD1-expressing MeA neurons that project to the BNST and ventromedial hypothalamus (VMH), regulating approach or avoidance of innate threats, respectively (Miller et al., 2019).

While previous literature has implicated the MeA in behavioral responses to unconditioned threats, our data suggests social defeat may prime the MeA to respond to a novel, non-aggressive conspecific as if they were a threat, leading to avoidance. This is in line with what has been demonstrated to some extent in rats (Luiten et al., 1985) and Syrian hamsters (Markham and Huhman, 2008). However, these studies reduced overall MeA activity either through physical lesion or pharmacological inactivation, and thus did not identify what neurochemicals may mediate this response. Dopamine in the MeA has been implicated to have a role in sexual behavior (Holder et al., 2015) and approach or avoidance of innate threats (Miller et al., 2019); however, to our knowledge, this study is the first to demonstrate that dopamine the MeA has a role in avoidance of conditioned stimuli that would normally not be perceived as threatening by a stress-naïve animal. The results of the present study suggest that defeat engineers dopamine-mediated activity that assesses social contact with unfamiliar conspecifics as “high threat” or “risky,” and elicits the same avoidance response that is innate to dangerous stimuli. Interestingly, in prairie voles, MeA and BNST cell bodies that project to the VMH increase in activity during exposure to pups, and pharmacological manipulation of the VMH significantly impairs approach and alloparental behaviors (Liu et al., 2019). Although the role of MeA DRD1 in this behavior was not examined, this finding, along with the discovery that MeA DRD1 projections to the BNST and VMH impact threat avoidance and social affiliation (Miller et al., 2019) and the results of the present study, may suggest a candidate neurocircuit for defeat-induced social behavior abnormalities. In addition, all these findings together indicate that the prairie vole’s unique social system could make this species an excellent model in examining the neurobiology behind defeat’s effects on various aspects of social behavior.

## CONCLUSIONS

Our model demonstrates equal defeat experience and subsequent behavioral phenotypes between male and female prairie voles. Thus, the prairie vole may provide a unique opportunity to make direct sex comparisons while investigating neurocircuitry responsible for defeat-induced behavioral abnormalities. In addition, we demonstrate a novel role of dopamine receptor activity in the MeA in approach/avoidance response following social defeat stress. With the recent emergence of transgenic prairie voles (Horie et al., 2018), this species may continue to rise as an attractive model in the study of neural mechanisms underlying stress-induced alterations in adaptive and maladaptive behavior. All together, these findings provide the prairie vole as a model of social defeat in both sexes, and implicate DRD1 in the MeA as a novel mechanism regulating avoidance of non-aggressive conspecifics following an aggressive encounter.

## Acknowledgements

We thank A. Swopes, B. Garcia, and M. Balay for assistance in data collection and behavioral analyses. We also thank Dr. S. Fowler for his assistance in quantifying vole behavior with the force-plate actometer in the open field test.

## REFERENCES

Arakawa, H., Arakawa, K., Deak, T., 2010. Oxytocin and vasopressin in the medial amygdala differentially modulate approach and avoidance behavior toward illness-related social odor. Neuroscience 171, 1141–1151.

Bjorkqvist, K., 2001. Social defeat as a stressor in humans. Physiol. Behav. 73, 435–442.

Bourke, C.H., Neigh, G.N., 2012. Exposure to repeated maternal aggression induces depressive-like behavior and increases startle in adult female rats. Behav. Brain Res. 227, 270–275.

Bowler, C.M., Cushing, B.S., Carter, C.S., 2002. Social factors regulate female-female aggression and affiliation in prairie voles. Physiol. Behav. 76, 559–566.

Burkett, J.P., Andari, E., Johnson, Z.V., Curry, D.C., de Waal, F.B., Young, L.J., 2016. Oxytocin-dependent consolation behavior in rodents. Science 351, 375–378.

Cadiz-Moretti, B., Otero-Garcia, M., Martinez-Garcia, F., Lanuza, E., 2016. Afferent projections to the different medial amygdala subdivisions: a retrograde tracing study in the mouse. Brain structure & function 221, 1033–1065.

Cao, J.L., Covington, H.E., 3rd, Friedman, A.K., Wilkinson, M.B., Walsh, J.J., Cooper, D.C., Nestler, E.J., Han, M.H., 2010. Mesolimbic dopamine neurons in the brain reward circuit mediate susceptibility to social defeat and antidepressant action. J. Neurosci. 30, 16453–16458.

Cervenka, S., Hedman, E., Ikoma, Y., Djurfeldt, D.R., Ruck, C., Halldin, C., Lindefors, N., 2012. Changes in dopamine D2-receptor binding are associated to symptom reduction after psychotherapy in social anxiety disorder. Translational psychiatry 2, e120.

Choi, G.B., Dong, H.-w., Murphy, A.J., Valenzuela, D.M., Yancopoulos, G.D., Swanson, L.W., Anderson, D.J., 2005. Lhx6 Delineates a Pathway Mediating Innate Reproductive Behaviors from the Amygdala to the Hypothalamus. Neuron 46, 647–660.

Dielenberg, R.A., Hunt, G.E., McGregor, I.S., 2001. ‘When a rat smells a cat’: the distribution of Fos immunoreactivity in rat brain following exposure to a predatory odor. Neuroscience 104, 1085–1097.

Fowler, S.C., Birkestrand, B.R., Chen, R., Moss, S.J., Vorontsova, E., Wang, G., Zarcone, T.J., 2001. A force-plate actometer for quantitating rodent behaviors: illustrative data on locomotion, rotation, spatial patterning, stereotypies, and tremor. J. Neurosci. Methods 107, 107–124.

Francis, T.C., Chandra, R., Friend, D.M., Finkel, E., Dayrit, G., Miranda, J., Brooks, J.M., Iniguez, S.D., O’Donnell, P., Kravitz, A., Lobo, M.K., 2015. Nucleus accumbens medium spiny neuron subtypes mediate depression-related outcomes to social defeat stress. Biol. Psychiatry 77, 212–222.

Gobrogge, K., Wang, Z., 2016. The ties that bond: neurochemistry of attachment in voles. Curr. Opin. Neurobiol. 38, 80–88.

Golden, S.A., Covington, H.E., 3rd, Berton, O., Russo, S.J., 2011. A standardized protocol for repeated social defeat stress in mice. Nat. Protoc. 6, 1183–1191.

Gray, C.L., Norvelle, A., Larkin, T., Huhman, K.L., 2015. Dopamine in the nucleus accumbens modulates the memory of social defeat in Syrian hamsters (Mesocricetus auratus). Behav. Brain Res. 286, 22–28.

Greenberg, G.D., Steinman, M.Q., Doig, I.E., Hao, R., Trainor, B.C., 2015. Effects of social defeat on dopamine neurons in the ventral tegmental area in male and female California mice. Eur. J. Neurosci. 42, 3081–3094.

Grippo, A.J., Cushing, B.S., Carter, C.S., 2007. Depression-like behavior and stressor-induced neuroendocrine activation in female prairie voles exposed to chronic social isolation. Psychosom. Med. 69, 149–157.

Harris, A.Z., Atsak, P., Bretton, Z.H., Holt, E.S., Alam, R., Morton, M.P., Abbas, A.I., Leonardo, E.D., Bolkan, S.S., Hen, R., Gordon, J.A., 2018. A Novel Method for Chronic Social Defeat Stress in Female Mice. Neuropsychopharmacology 43, 1276–1283.

Holder, M.K., Veichweg, S.S., Mong, J.A., 2015. Methamphetamine-enhanced female sexual motivation is dependent on dopamine and progesterone signaling in the medial amygdala. Horm. Behav. 67, 1–11.

Horie, K., Inoue, K., Suzuki, S., Adachi, S., Yada, S., Hirayama, T., Hidema, S., Young, L.J., Nishimori, K., 2018. Oxytocin receptor knockout prairie voles generated by CRISPR/Cas9 editing show reduced preference for social novelty and exaggerated repetitive behaviors. Horm. Behav.

Huhman, K.L., Solomon, M.B., Janicki, M., Harmon, A.C., Lin, S.M., Israel, J.E., Jasnow, A.M., 2003. Conditioned defeat in male and female Syrian hamsters. Horm. Behav. 44, 293–299.

Kendler, K.S., Hettema, J.M., Butera, F., Gardner, C.O., Prescott, C.A., 2003. Life event dimensions of loss, humiliation, entrapment, and danger in the prediction of onsets of major depression and generalized anxiety. Arch. Gen. Psychiatry 60, 789–796.

Kenkel, W.M., Suboc, G., Carter, C.S., 2014. Autonomic, behavioral and neuroendocrine correlates of paternal behavior in male prairie voles. Physiol. Behav. 128, 252–259.

Li, C.I., Maglinao, T.L., Takahashi, L.K., 2004. Medial amygdala modulation of predator odor-induced unconditioned fear in the rat. Behav. Neurosci. 118, 324–332.

Li, L.F., Yuan, W., He, Z.X., Wang, L.M., Jing, X.Y., Zhang, J., Yang, Y., Guo, Q.Q., Zhang, X.N., Cai, W.Q., Hou, W.J., Jia, R., Tai, F.D., 2018. Involvement of oxytocin and GABA in consolation behavior elicited by socially defeated individuals in mandarin voles. Psychoneuroendocrinology 103, 14–24.

Lieberwirth, C., Liu, Y., Jia, X., Wang, Z., 2012. Social isolation impairs adult neurogenesis in the limbic system and alters behaviors in female prairie voles. Horm. Behav. 62, 357–366.

Liu, Y., Donovan, M., Jia, X., Wang, Z., 2019. The ventromedial hypothalamic circuitry and male alloparental behavior in a socially monogamous rodent species. Eur. J. Neurosci.

Liu, Y., Young, K.A., Curtis, J.T., Aragona, B.J., Wang, Z., 2011. Social Bonding Decreases the Rewarding Properties of Amphetamine through a Dopamine D1 Receptor-Mediated Mechanism. The Journal of Neuroscience 31, 7960–7966.

Luiten, P.G.M., Koolhaas, J.M., de Boer, S., Koopmans, S.J., 1985. The cortico-medial amygdala in the central nervous system organization of agonistic behavior. Brain Res. 332, 283–297.

Markham, C.M., Huhman, K.L., 2008. Is the medial amygdala part of the neural circuit modulating conditioned defeat in Syrian hamsters? Learn. Memory 15, 6–12.

McGregor, I.S., Hargreaves, G.A., Apfelbach, R., Hunt, G.E., 2004. Neural Correlates of Cat Odor-Induced Anxiety in Rats: Region-Specific Effects of the Benzodiazepine Midazolam. The Journal of Neuroscience 24, 4134–4144.

McNeal, N., Appleton, K.M., Johnson, A.K., Scotti, M.L., Wardwell, J., Murphy, R., Bishop, C., Knecht, A., Grippo, A.J., 2017. The protective effects of social bonding on behavioral and pituitary-adrenal axis reactivity to chronic mild stress in prairie voles. Stress 20, 175–182.

Meredith, M., Westberry, J.M., 2004. Distinctive Responses in the Medial Amygdala to Same-Species and Different-Species Pheromones. The Journal of Neuroscience 24, 5719–5725.

Miller, S.M., Marcotulli, D., Shen, A., Zweifel, L.S., 2019. Divergent medial amygdala projections regulate approach–avoidance conflict behavior. Nat. Neurosci.

Müller, M., Fendt, M., 2006. Temporary inactivation of the medial and basolateral amygdala differentially affects TMT-induced fear behavior in rats. Behav. Brain Res. 167, 57–62.

Northcutt, K.V., Lonstein, J.S., 2009. Social contact elicits immediate-early gene expression in dopaminergic cells of the male prairie vole extended olfactory amygdala. Neuroscience 163, 9–22.

O’Connell, L.A., Hofmann, H.A., 2011. Genes, hormones, and circuits: an integrative approach to study the evolution of social behavior. Front. Neuroendocrinol. 32, 320–335.

Pardo-Bellver, C., Cadiz-Moretti, B., Novejarque, A., Martinez-Garcia, F., Lanuza, E., 2012. Differential efferent projections of the anterior, posteroventral, and posterodorsal subdivisions of the medial amygdala in mice. Front. Neuroanat. 6.

Reguilon, M.D., Montagud-Romero, S., Ferrer-Perez, C., Roger-Sanchez, C., Aguilar, M.A., Minarro, J., Rodriguez-Arias, M., 2017. Dopamine D2 receptors mediate the increase in reinstatement of the conditioned rewarding effects of cocaine induced by acute social defeat. Eur. J. Pharmacol. 799, 48–57.

Silk, J., Cheney, D., Seyfarth, R., 2013. A practical guide to the study of social relationships. Evolutionary anthropology 22, 213–225.

Smith, A.S., Lieberwirth, C., Wang, Z., 2013. Behavioral and physiological responses of female prairie voles (Microtus ochrogaster) to various stressful conditions. Stress 16, 531–539.

Smith, A.S., Wang, Z., 2014. Hypothalamic oxytocin mediates social buffering of the stress response. Biol. Psychiatry 76, 281–288.

Takahashi, A., Chung, J.R., Zhang, S., Zhang, H., Grossman, Y., Aleyasin, H., Flanigan, M.E., Pfau, M.L., Menard, C., Dumitriu, D., Hodes, G.E., McEwen, B.S., Nestler, E.J., Han, M.H., Russo, S.J., 2017. Establishment of a repeated social defeat stress model in female mice. Sci. Rep. 7, 12838.

Taymans, S.E., DeVries, A.C., DeVries, M.B., Nelson, R.J., Friedman, T.C., Castro, M., Detera-Wadleigh, S., Carter, C.S., Chrousos, G.P., 1997. The hypothalamic-pituitary-adrenal axis of prairie voles (Microtus ochrogaster): evidence for target tissue glucocorticoid resistance. Gen. Comp. Endocrinol. 106, 48–61.

Thacker, J.S., Yeung, D.H., Staines, W.R., Mielke, J.G., 2016. Total protein or high-abundance protein: Which offers the best loading control for Western blotting? Anal. Biochem. 496, 76–78.

Toth, I., Neumann, I.D., 2013. Animal models of social avoidance and social fear. Cell Tissue Res. 354, 107–118.

Trainor, B.C., Pride, M.C., Villalon Landeros, R., Knoblauch, N.W., Takahashi, E.Y., Silva, A.L., Crean, K.K., 2011. Sex Differences in Social Interaction Behavior Following Social Defeat Stress in the Monogamous California Mouse (Peromyscus californicus). PLoS One 6, e17405.

Trainor, B.C., Takahashi, E.Y., Campi, K.L., Florez, S.A., Greenberg, G.D., Laman-Maharg, A., Laredo, S.A., Orr, V.N., Silva, A.L., Steinman, M.Q., 2013. Sex differences in stress-induced social withdrawal: Independence from adult gonadal hormones and inhibition of female phenotype by corncob bedding. Horm. Behav. 63, 543–550.

Wang, L., Zhu, Z., Hou, W., Zhang, X., He, Z., Yuan, W., Yang, Y., Zhang, S., Jia, R., Tai, F., 2018. Serotonin signaling trough prelimbic 5-HT1A receptors modulates CSDS induces behavioural changes in adult female voles. Int. J. Neuropsychopharmacol.

Willner, P., Hale, A.S., Argyropoulos, S., 2005. Dopaminergic mechanism of antidepressant action in depressed patients. J. Affect. Disord. 86, 37–45.

