## Supplementary Information for "Regulation of defeat-induced social avoidance by medial amygdala DRD1 in male and female prairie voles"

**Figure S1:** ***Experimental timelines.*** **A)** Timeline for Experiment 1, with one animal receiving control, single, or repeated defeat conditioning. **B)** Timeline for Experiment 2, with both cagemates receiving either control or repeated defeat conditioning. **C)** Timeline for Experiment 3, with both cagemates receiving control or repeated defeat conditioning, then one of two drug treatments 30 minutes prior to SPA (CSF or SKF 38393 for controls, CSF or SCH 23390 for defeated voles). **D)** Timeframe between drug administration and SPA testing. Animals had a total 30 minute drug activation time prior to beginning the SPA test. *SPA- Social preference/avoidance test; EPM- Elevated plus maze test; FS-Forced swim test; OFT- Open field test; SPT- Sucrose preference test.*

### Figure S1

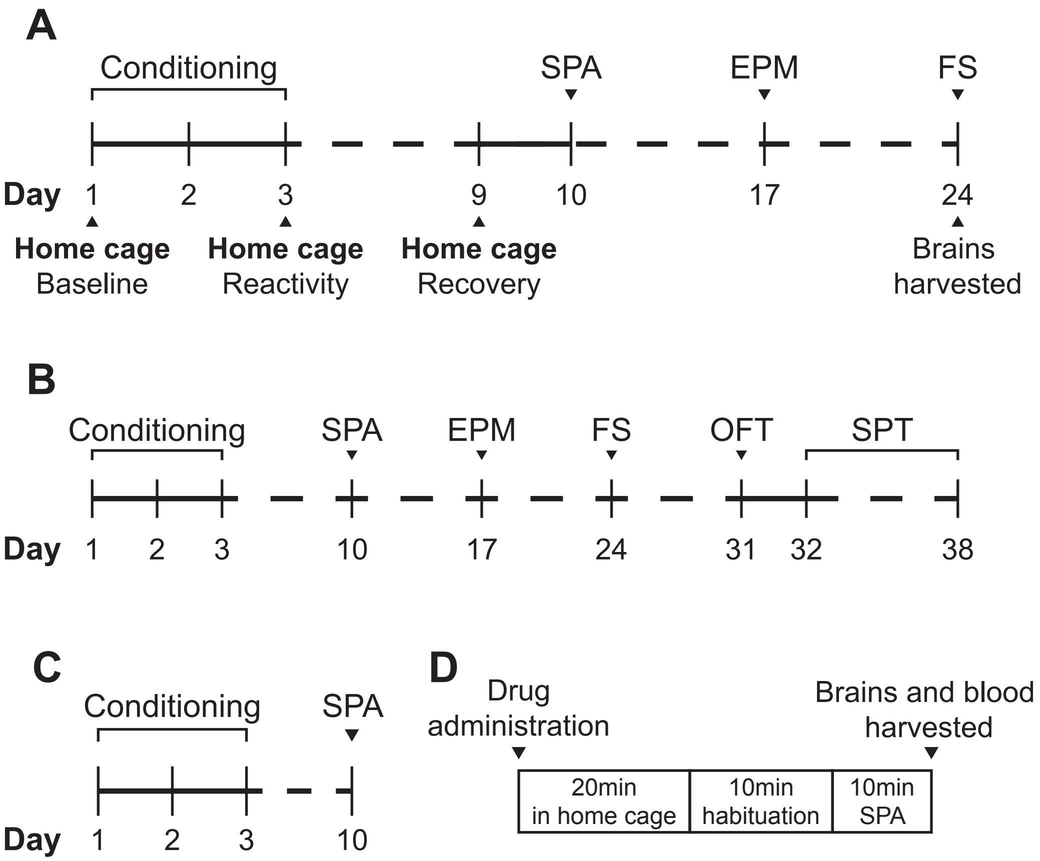
A
Conditioning
SPA
EPM
FS
Day 1 2 3 9 10 17 24
Home cage
Baseline
Home cage
Reactivity
Home cage
Recovery
Brains
harvested
B
Conditioning
SPA
EPM
FS
OFT
SPT
Day 1 2 3 10 17 24 31 32 38
C
Conditioning
SPA
Day 1 2 3 10
D
Drug
administration
Brains and blood
harvested
20min
in home cage
10min
habituation
10min
SPA

**Table S1: Ethogram of behaviors analyzed during defeat.**

| **Offensive Behaviors** | **Definition** |
|---|---|
| *Attack* | Attacker initiates strike/rough and tumble/boxing |
| *Chase* | Attacker running after opponent |
| *Lateral Threat* | Attacker directs its flank toward the opponent |
| *Upright posture* | Rearing upright with forelimbs extended |
| *Stamp* | Attacker stomps the ground or kicks up bedding |
| *Lunge* | Mock lunge/lunge with body but no attack results |
| **Defensive behaviors** | **Definition** |
| *Retaliatory Attack* | Opponent attacks immediately after attacker attack |
| *Leap Escape* | Jumping after a lunge/attack/or toward wall |
| *Flee* | Opponent runs after attack/threat from attacker |
| *Supine* | Opponent lays on back after an attack/threat |
| *Defensive posture* | Lean back upright or sitting (forepaw may be up) |

**Table S2: Antibodies used for western blot anaylsis of DRD1 and DRD2 expression.** Antibody substitutions were due to manufacturer lot replacement and were individually validated before use. *DRD1- Dopamine receptor 1; DRD2- Dopamine receptor 2; PFC- Prefrontal cortex; NAcc- Nucleus accumbens; BLA- Basolateral amygdala; CeA- Central amygdala; MeA- Medial amygdala.*

| Antigen | Regions | Manufacturer | Cat # | Concentration |
|---|---|---|---|---|
| DRD1 | NAcc | Santa Cruz | sc-336600 | 1:500 |
| DRD1 | CeA, MeA, BLA, PFC | Abcam | ab20066 | 1:4000 |
| DRD2 | NAcc, CeA, MeA | Santa Cruz | sc-5303 | 1:500 |
| DRD2 | BLA, PFC | EMD Millipore | ab5084P | 1:1000 |
| Mouse IgG | all | Santa Cruz | sc-516102 | 1:10000 |
| Rabbit IgG | all | Santa Cruz | sc-2357 | 1:10000 |

**Table S3:** ***Breakdown of statistics for individual behaviors observed during defeat conditioning.***

***Offensive Behaviors (frequency)***

| | Aggressor | | Subject | | Stats | | |
|---|---|---|---|---|---|---|---|
| **Behavior** | **Male** | **Female** | **Male** | **Female** | **Effect** | **F(1,14)** | **p** |
| Attack | 14.667±1.622 | 9.500±0.807 | 0.250±0.083 | 0.167±0.089 | Social status x sex | 7.764 | 0.015 |
| Chase | 17.000±1.790 | 11.792±2.095 | 0.333±0.154 | 0.292±0.098 | Social status | 99.121 | 0.000 |
| Lateral Threat | 2.542±0.831 | 1.375±0.596 | 0.417±0.280 | 0.167±0.089 | Social status | 10.337 | 0.006 |
| Upright posture | 0.417±0.122 | 0.958±0.364 | 1.750±0.519 | 1.375±0.572 | N/A | 3.751 | *n.s.* |
| Stamp | 0.833±0.227 | 2.125±0.743 | 0.000±0.000 | 0.042±0.042 | Social status | 13.676 | 0.002 |
| Lunge | 0.765±0.270 | 0.309±0.109 | 2.625±0.592 | 2.500±0.748 | Social status | 21.000 | 0.000 |

***Defensive Behaviors (frequency)***

| | Aggressor | | Subject | | Stats | | |
|---|---|---|---|---|---|---|---|
| **Behavior** | **Male** | **Female** | **Male** | **Female** | **Effect** | **F (1,14)** | **p** |
| Retaliatory Attack | 0.125±0.044 | 0.208±0.074 | 1.458±0.516 | 0.250±0.088 | Social status x sex | 4.921 | 0.044 |
| Leap Escape | 0.458±0.281 | 0.083±0.055 | 10.417±2.016 | 8.875±2.631 | Social status | 34.109 | 0.000 |
| Flee | 1.000±0.383 | 1.000±0.295 | 24.333±2.720 | 20.042±3.545 | Social status | 90.204 | 0.000 |
| Supine | 0.125±0.088 | 0.083±0.083 | 3.917±1.467 | 2.000±0.639 | Social status | 13.855 | 0.002 |
| Defensive posture | 0.625±0.248 | 1.083±0.503 | 17.042±1.487 | 13.958±0.878 | Social status | 329.755 | 0.000 |
